# Functional Specialization and Distributed Processing across Marmoset Lateral Prefrontal Subregions

**DOI:** 10.1101/2024.02.29.582669

**Authors:** Raymond Ka Wong, Janahan Selvanayagam, Kevin D Johnston, Stefan Everling

## Abstract

A prominent aspect of the organization of primate lateral prefrontal cortex (lPFC) is its division into a number of cytoarchitecturally distinct subregions. Investigations in macaque lPFC using neurophysiological approaches have provided much insight into the functions associated with these subregions; however, our understanding is based largely on a patchwork of findings from many studies and across many animals, rarely covering the entire lPFC in individual subjects. Here, we leveraged the small size and lissencephalic cortex of the common marmoset (*Callithrix jacchus*) to characterize the responses of large numbers of single lPFC neurons to a diverse collection of test stimuli recorded across sets of lPFC subregions using high-density microelectrode arrays. Untethered extracellular electrophysiological recordings were obtained from two adult marmosets with 4 x 4 mm 96-channel Utah arrays implanted in lPFC, covering areas 8aD, 8aV, 9, 10, 46D, 46V and 47. We employed a test battery comprised of a variety of visual stimuli including faces and body parts, auditory stimuli including marmoset calls, and a spatial working memory task. Task-modulated units and units responsive to different stimulus modalities were distributed throughout the lPFC. Visual, auditory and call-selective units were distributed across all lPFC subregions. Neurons with contralateral visual receptive fields were found in 8aV and 8aD. Neurons responsive to faces and saccade-related units were found in 8aV, 8aD, 10, 46V and 47. These findings demonstrate that responses to some stimuli are relatively restricted within specific lPFC subregions, while others are more distributed throughout the marmoset lPFC.

## Introduction

The highly evolved lateral prefrontal cortex (lPFC) in primates is involved in higher-order cognitive processes, including mental representation of abstract rules (Wallis et al. 2001; Everling and DeSouza 2005; Arnsten et al. 2012), working memory (Fuster and Alexander 1971), executive control (Desimone and Duncan 1995; Miller and Cohen 2001), as well as responses to different stimulus modalities (visual stimuli, auditory, including complex stimuli such as faces and conspecific calls (O Scalaidhe et al. 1999; Romanski and Goldman-Rakic 2002; Sugihara et al. 2006; Romanski and Averbeck 2009; Riley et al. 2016; Haile et al. 2019). Studies of anatomical connectivity and cytoarchitecture show that the PFC is partitioned into numerous major cytoarchitectonic regions and even finer subdivisions by some parcellations (Barbas and Pandya 1989; Petrides 2005; Averbeck and Lee 2006; Sallet et al. 2013). In addition to the anatomical connectivity, O’Reilly (2010) argues that there is a systematic functional organization across lPFC areas, as functional architecture is a canonical property of the cerebral cortex (Van Essen and Glasser 2018). Addressing the functional parcellation of the lPFC is critical to our understanding of how the lPFC implements executive functions. As noted above, investigations in macaque lPFC have provided much insight into area-specific functionality, but our understanding is based primarily on area-specific parcellation and recordings across multiple animals and never spanning the entire lPFC in one animal. This potentially obscures some functional differences as varying tasks and training protocols have been shown to affect both stimulus representations and their distribution within lPFC (Bichot et al. 1996; Rao et al. 1997).

A nonhuman primate species offering practical advantages for large-scale mapping of cortical areas, is the small New World common marmoset (*Callithrix jacchus*). This species’ relatively lissencephalic cortex also offers the opportunity for laminar electrophysiological recordings and optical imaging (Sadakane et al. 2015; Kondo et al. 2018; Johnston et al. 2019). Consequently, considerable effort has been directed toward an understanding of the cytoarchitectural differentiation of PFC subregions, and indeed the marmoset lPFC is comprised of a number of cytoarchitecturally distinct subfields believed to be homologous with those of macaques and humans (Burman et al. 2006; Reser et al. 2013). Owing to the relatively recent surge in popularity of the marmoset model however (Mitchell and Leopold 2015; Okano 2021), the functions associated with these subregions are relatively poorly understood in this species and are an area of intensive investigation (Blum et al. 1982; Hung et al. 2015; Johnston et al. 2019; Liu et al. 2019; Schaeffer, Gilbert, Gati, et al. 2019; Schaeffer, Gilbert, Hori, et al. 2019; Selvanayagam et al. 2019; Schaeffer et al. 2020; Feizpour et al. 2021; Jovanovic et al. 2022; Wong et al. 2023). Given the well-established link between the lPFC and cognition, and the unique potential of the marmoset model for deriving an understanding of the cortical microcircuitry underlying aspects of social cognition such as vocal communication (Miller et al. 2016; Jovanovic et al. 2022; Samandra et al. 2022; Grijseels et al. 2023), establishing a correspondence between the structural and functional organization of lPFC with respect to relatively simple tasks and social stimuli of multiple modalities is needed to provide an empirical foundation for interpretation of these more complex processes.

Here, we sought to characterize the response properties of single neurons in lPFC subregions using electrophysiological single neuron recordings spanning a large portion of the lPFC within individual animals. Wireless extracellular electrophysiological recordings were obtained using a data-logging recording system from two adult marmosets with a 96-channel Utah array (4×4 mm, 1.5 mm electrode length, 400μm pitch) implanted in the left lPFC, covering areas 8aV, 8aD, 9, 10, 46D, 46V and 47. To characterize these lPFC subregions, we recorded neural activity in response to a variety of visual and auditory stimuli and during the performance of a spatial working memory task.

## Methods

### Subjects

Data was collected from two adult female common marmosets (*Callithrix jacchus*; Marmoset A, 26 months; Marmoset B, 24 months). All experimental procedures conducted were in accordance with the Canadian Council of Animal Care policy on the care and use of laboratory animals and a protocol approved by the Animal Care Committee of the University of Western Ontario Council on Animal Care. The animals were under the close supervision of university veterinarians.

### Array surgery

Animals underwent an aseptic surgical procedure under general anesthesia in which 96-channel electrode arrays (4 mm x 4 mm; 1.5 mm electrode length; 400 µm pitch; iridium oxide tips) (Blackrock Neurotech, Salt Lake City, US) were implanted in the left PFC (see Selvanayagam et al. 2019 for details). During this surgery, a microdrill was used to perform a ∼5 mm craniotomy which was enlarged as necessary using a rongeur. The dura was removed, and the array was manually inserted into the lateral PFC; wires and connectors were fixed to the skull using dental adhesive and resin cement (All-Bond Universal and Duo-Link, Bisco Dental Products). Once implanted, the array site was covered with a thin layer of silicone adhesive (Kwik Sil; World Precision Instruments). A screw hole was drilled into the right side of the skull to place a stainless-steel ground screw. The ground wire of the array was then tightly wound around the base of the screw to ensure a stable electrical connection. A combination recording chamber/head holder (Johnston et al. 2018) was placed around the array and connectors and fixed in place using further layers of dental adhesive and resin cement. Finally, a removable protective cap was placed on the chamber to protect the 3×32 channel Omnetics connector.

### Neural recordings

After recovery from array implantation, we verified that electrode contacts were within the cortex by monitoring extracellular neural activity using the SpikeGadgets’ data acquisition system (SpikeGadgets, San Francisco, US). Upon observing single- or multiunit activity at multiple sites of the array for approximately three weeks, we commenced unrestrained datalogger recordings of extracellular activity from the 96 implanted electrodes. A detailed description of these unrestrained datalogger-based recordings is presented in Wong et al. (2023). Initially, neural data underwent processing with a common median filter to mitigate large movement-related artifacts. Subsequently, the data were further processed using a 4-pole Butterworth high-pass filter with a cutoff frequency of 500 Hz. Spike detection and sorting were then carried out offline using Plexon Offline Sorter v3. For our analysis, we included only those clearly isolated single units that exhibited baseline discharge rates exceeding 0.5 Hz. Specifically, we analyzed 2574 units from marmoset B and 2188 units from marmoset A.

### Visual and Auditory Stimulus Presentation and Eye Movement Monitoring

For all visual receptive field mapping, visual stimulus presentation, and some auditory stimulus presentation sessions, we recorded neural activity while animals were head restrained. In these sessions, marmosets were seated in a custom-designed primate chair (Johnston et al. 2018) inside a sound attenuating chamber (Crist Instrument Co. Hagerstown MD), with the head restrained. A spout was placed at animals’ mouth to allow delivery of a viscous liquid reward (acacia gum) via an infusion pump (Model NE-510, New Era Pump Systems, Inc., Farmingdale, New York, USA). All visual stimuli were presented on a CRT monitor (ViewSonic Optiquest Q115, 76 Hz non-interlaced, 1600 x 1280 resolution) using Monkeylogic (Hwang et al. 2019) on an ASUS UX430U Notebook PC running Windows 10. Eye positions were digitally recorded at 1 kHz via infrared video tracking of the left pupil (EyeLink 1000, SR Research, Ottawa, ON, Canada). Auditory stimulus presentation was controlled by Raspberry Pi 3 Model B and presented on a Bose Soundlink III speaker (Bose Corporation, Framingham Mass.) connected to the audio output of the raspberry pi and placed at a distance of 10cm centred in front of the animals, and 12cm below head level.

### Experimental Design and Stimulus Presentation

#### Delayed-match-to-location (DML) task

Marmosets performed a DML task on an in-house developed touchscreen testing box attached to the home cage (for details of touchscreen training protocol, see (Wong et al. 2023). Each trial began with the presentation of a sample stimulus (filled blue or pink circle, 3 cm diameter) on a grey background at one of the four corner locations of the touchscreen display for a duration of 2.5 s. This was followed by a 2s delay period in which the screen remained blank. After the delay period, choice stimuli (filled blue or pink circles, 3 cm diameter) were presented at each of the four corner locations and the animal was required to touch the location matching the previously presented stimulus to obtain a liquid reward of 0.075-0.1 mL 50/50 mix of 1:1 acacia gum powder and water with liquid marshmallow. The reward was delivered via an infusion pump (model NE-510; New Era Pump Systems) through a liquid spout placed in front of the touchscreen monitor (Elo 1002L). Trials were separated by 5 s intertrial periods.

#### Visual Receptive Field Mapping

To map visual receptive fields in the lPFC of marmosets, we conducted a series of trials where a pseudorandom sequence of visual stimuli was displayed on the monitor. Each trial commenced with the animal fixating on a central dot for 500 ms. Following this initiation, circular stimuli, each subtending 0.2°, were presented rapidly at 9 pseudorandom locations selected from a pool of 48 possible sites (arranged in a 7×7 grid, covering +/-12 degrees along both the ordinate and abscissa). Although eye position was monitored, animals were not required to maintain fixation during stimulus presentation. Stimulus onset asynchrony was set at 300 ms, with an interstimulus interval of 100 ms and an intertrial interval of 1-2 seconds. To sustain alertness of the animals, a liquid reward was dispensed at the end of each trial.

#### Presentation of Visual Stimuli

To examine lPFC neuron responses to complex visual stimuli, we presented one of two distinct image sets in each recording session: 1) a collection of human and marmoset faces, scrambled faces, and objects, and 2) images featuring arms, bodies, and faces of marmosets. In set 1, there were 6 human faces, 11 marmoset faces, 23 objects, and their corresponding scrambled versions (each stimulus presented roughly 19 times per session). The scrambling of faces involved dividing them into spatially-segmented blocks and randomly shuffling these within and then across rows. Set 2 comprised 24 stimuli of each type (each presented about 14 times per session). All images subtended 3.5 degrees of the visual angle. During trials, the animals needed to fixate on a central dot (0.4°) for 300-500 ms before the display of a single image for 250 ms on the monitor. An inter-trial interval of 2 seconds was maintained. Trials were aborted if the animal’s gaze strayed beyond a 9-degree radius window centered on the displayed image

#### Presentation of Auditory Stimuli

Acoustic stimuli of different conspecific calls (phee, trill, trillphee, twitter, chirp, ek, tsik, chatter) and scrambled (shuffled 25 ms windows over a 250 ms radius) versions of these calls were presented in three different contexts (head-restrained in a primate chair, head-unrestrained in a primate chair, and freely moving in their home cage) during each recording session. The order of context presentation was counterbalanced. In each context, the presentation of calls was pseudo-randomized with a 1-1.5 s inter-trial-interval. Stimuli were obtained in-house and from an open-source database (Landman et al. 2020). Our dataset consisted of eight different call types and two different calls for each call type. On average, each call was presented 48 times in each context and session. Calls were recorded in-house or obtained from an open source database (Landman et al. 2020).

### Data analysis

Analysis was performed using custom code written in Matlab (MathWorks) and python. Statistical significance was evaluated at an alpha level of p < .02.

#### DML task

Previously published data from Wong and colleagues (2023) was included here. Activity within distinct non-overlapping task epochs was assessed using analyses of variance (ANOVA) to compare the mean discharge rates of each neuron for each condition (four spatial locations) and each task epoch (baseline, sample, delay, pre-response and post-response). The baseline epoch was defined as the 1.5 s prior to the onset of the sample stimulus to the time of sample stimulus presentation. The sample epoch was defined as the 100 ms after the onset of the sample stimulus to the time of stimulus offset (2.4 s). We excluded the first 100 ms to avoid any potential sluggish sample-related activity contaminating estimates of delay-related activity. The delay epoch was 2 s in duration. The pre-response epoch was defined as the 300 ms period prior to the touch response and the post-response epoch was defined as the 1000 ms immediately after the touch response.

#### Visual Receptive Field Mapping

Analyses of variance (ANOVAs) were used to compare the mean discharge rates of each neuron for each task epoch (baseline, post-stimulus) and condition (ipsilateral, contralateral). The baseline epoch was defined as the 200 ms prior to the trial onset for all stimulus presentations in that trial. The post-stimulus epoch was 50-150 ms after stimulus onset. We excluded all stimulus presentations in which a saccade was made within +/-200ms of stimulus onset.

#### Categorical Visual Stimuli

ANOVAs were used to compare the mean discharge rates of each neuron for each epoch (baseline, post-stimulus) and condition (human face, marmoset face, scrambled faces, objects). The baseline epoch was defined as the 450 ms prior to the stimulus onset to 50 ms after stimulus onset. The peak response of a given neuron and its response to categorical images varied post-stimulus presentation. To determine the peak response time, we calculated an average spike density function with a kernel filter that resembles a postsynaptic potential (Thompson et al. 1996) and determined the maximum peak or trough. Time constants for the growth and decay phases were set at 1 ms and 20 ms, respectively. The start and end of the post-stimulus epoch for each neuron was then defined as the time at which the discharge rates reached 70% of the maximum response. A selectivity index for category and individual images were calculated as a contrast ratio using mean firing rates (FR):

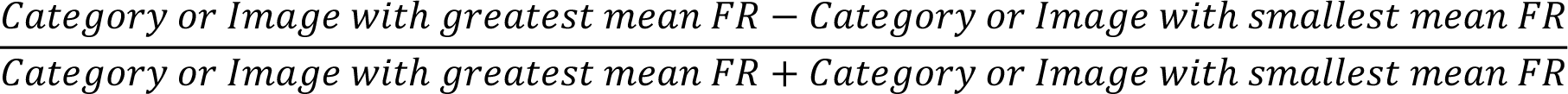

#### Saccades

In sessions in which we presented visual stimuli of different categories, we analysed the saccades made within each session. Saccades that had an amplitude less than 0.5 and greater than 20 visual degrees, as well as velocity greater than 1200 degree/s were excluded. ANOVAs were used to compare the mean discharge rates of each neuron for each epoch (baseline, pre-saccadic, peri-saccadic, post-saccadic). The baseline epoch was defined as the 200-100 ms prior to saccade onset. The pre-saccadic epoch was 100-25 ms prior to saccade onset. The peri-saccadic epoch was 25 ms prior to saccade onset to 25 ms after saccade end. The post-saccadic epoch was 25-100 ms after saccade end.

#### Auditory Responses

ANOVAs were used to compare the mean discharge rates of each neuron for each epoch (baseline, post-stimulus), call type (phee, trill, trillphee, twitter, chirp, ek, tsik, chatter, scrambled) and context (head-restrained and head-unrestrained in a primate chair, freely moving in their home cage). The baseline epoch was defined as the 200 ms prior to the stimulus onset to 50 ms after stimulus onset. We noted that most neurons had an onset response to auditory stimuli which occurred at varying latencies and that the decay of this response varied across neurons and call types. To account for these variations in timing and response dynamics, accurately capture statistically these responses, and effectively allow a comparison between different call types, we constructed dynamic response epochs by aligning activity on stimulus onset and determining the times at which the observed post-stimulus peak in activity reached 70% of the maximum discharge rate on both the rising and falling phases of the response. To determine the peak response time, we calculated an average spike density function with a kernel filter resembling a postsynaptic potential (Thompson et al. 1996) and found the maximum peak or trough. Time constants for the growth and decay phases were set at 1 ms and 20 ms, respectively. We then computed the mean discharge rate within this epoch, the beginning and end of which were delineated by these times. In practice these resulting post-stimulus epochs had a range of durations from 25-125 ms.

### Natural grouping of neural responses

We used an unsupervised clustering algorithm described by Kiani and colleague (2015) to reveal potential spatially segregated clusters within the lPFC. For each recording session, we identified natural physiological groupings of recorded units based on the dissimilarity of their responses (Kiani et al. 2015). The response dissimilarity reflects covariation of neural responses and can take any value between 0 (perfect correlation) and 2 (perfect anti-correlation). We defined the neural response vector for each unit in 30 ms non-overlapping bins from the beginning to end of recording sessions, independent of task epochs and the animal’s behaviour. Dissimilarities for all possible pairs of units in a given session was calculated and a 96 x 96 dissimilarity matrix for all possible pairs of recording channels was created for each session. Dissimilarity matrixes across sessions were averaged for each animal separately.

We applied nonlinear multi-dimensional scaling (MDS) to the averaged dissimilarity matrix to create a low dimensional representation that retained the pairwise relationships as much as possible. To explore the spatial relationship between recording channels, we chose a unique colour for each recording channel based on its location in the 2-dimensional MDS map and a 2D colour map. Locations on the array with similar colours represent natural physiological groupings of recorded units.

### Array localization

Marmosets were euthanized at the end of the data acquisition process to prepare the brains for ex-vivo MRI scans (Wong et al. 2023). The animals were deeply anesthetized with 20 mg/kg of ketamine plus 0.025 mg/kg medetomidine and 5% isoflurane in 1.4–2% oxygen to reach beyond the surgical plane (i.e., no response to toe pinching or cornea touching). They were then transcardially perfused with 0.9% sodium chloride irrigation solution, followed by 10% buffered formalin. The brain was then extracted and stored in 10% buffered formalin for more than a week before ex-vivo magnetic resonance imaging (MRI). On the day of the scan, the brain was transferred to another container for imaging and immersed in a fluorine-based lubricant (Christo-lube; Lubrication Technology) to improve homogeneity and avoid susceptibility artifacts at the boundaries. Ex-vivo MRI was performed on a 9.4T 31 cm horizontal bore magnet (Varian/Agilent, Yarnton, UK) and Bruker BioSpec Avance III console with the software package Paravision-7 (Bruker BioSpin Corp, Billerica, MA), a custom-built high-performance 15-cm-diameter gradient coil with 400 mT/m maximum gradient strength (xMR, London, CAN; Peterson et al. 2018), and an mp30 (Varian Inc., Palo Alto, USA) transmit/receive coil. High resolution (100×100×100 µm) T2-weighted images were acquired for each animal. The raw MRI images were converted to NifTI format using dcm2niix (Li et al. 2016) and the MRIs were nonlinearly registered to the ultra-high-resolution ex-vivo NIH template brain (Liu et al. 2018), that contains the location of cytoarchitectonic boundaries of the Paxinos atlas (Paxinos et al. 2012), using Advanced Normalization Tools (ANTs; Avants et al. 2011) software. The resultant transformation matrices were then applied to the cytoarchitectonic boundary image included with the NIH template brain atlas. These cytoarchitectonic boundaries overlayed on the registered ex-vivo anatomical T2 images were used to reconstruct the location of the implanted array in each marmoset (Supplementary Figure 1).

## Results

Overall, neurons across many areas of marmoset PFC exhibited significant modulations in activity during a cognitive task as well as during presentations of visual and auditory stimuli and across differing experimental contexts. Single neuron examples depicting these modulations are presented in Figure 1. As noted in our previously published work (Wong et al. 2023) we observed task-related activity during the sample, delay, and response epochs of the DML task, with many neurons exhibiting modulations in one or more of these epochs (Figure 1A). We noted that a range of visual stimuli including faces, objects, and body parts also evoked robust responses which were selective for the stimulus type in many cases (Figure 1B). We additionally observed pre-saccadic and robust post-saccadic activity (Figure 1C). Finally, we observed call-selective auditory responses in PFC neurons (Figure 1D) that were in some cases modulated across experimental contexts including head-restrained, head-unrestrained, and within the home cage (Figure 1D). Specific analyses investigating these responses are detailed below.

**Figure 1.**
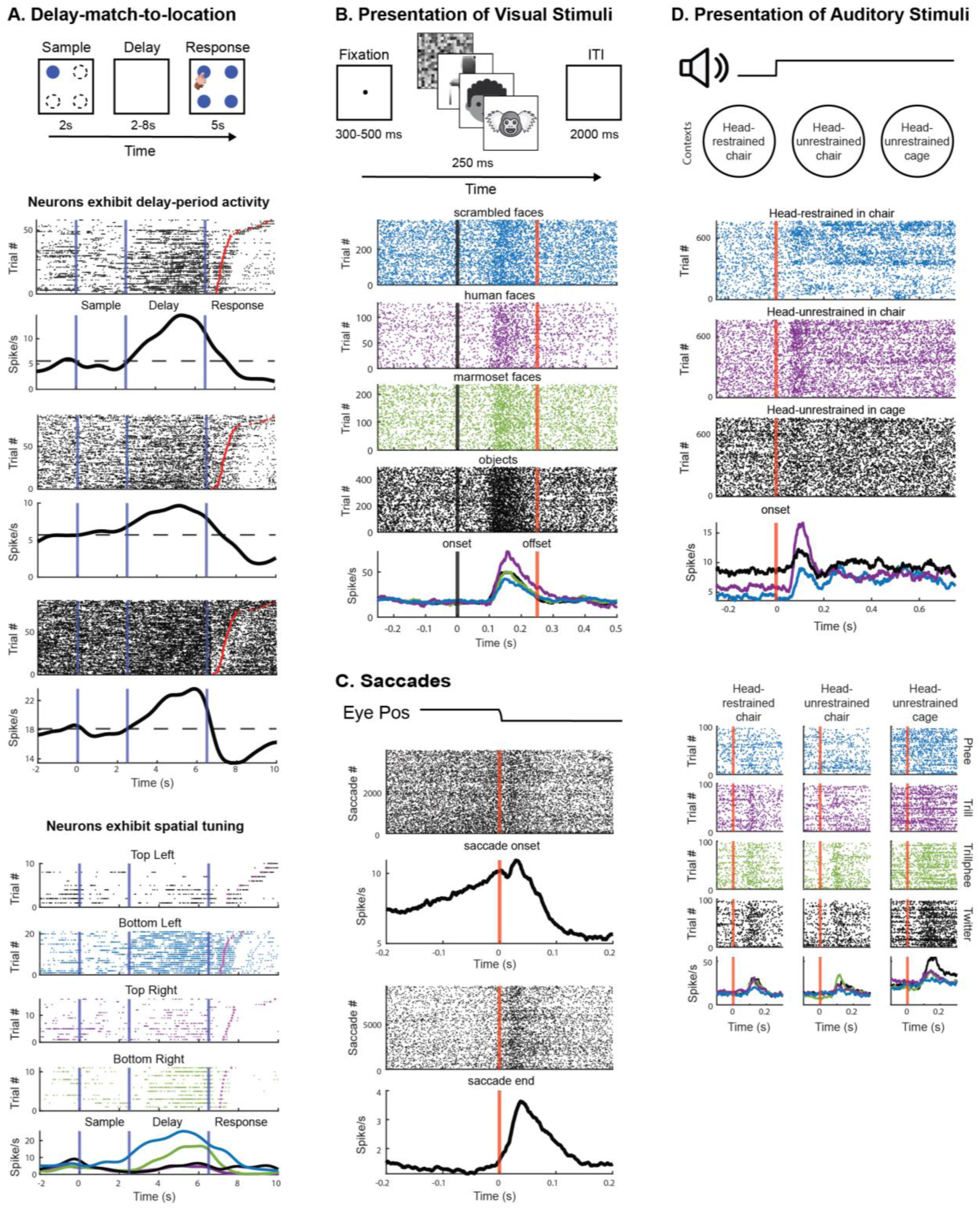
Example single Marmoset lPFC neurons modulated by task and stimulus. A) Top panel, task timeline. Rasters and spike density functions in panels below depict single units exhibiting significant modulations in discharge rate during the delay epoch. Panels below depict a single neuron exhibiting spatial tuning. B) Top panel, task timeline. Below depicts a single neuron exhibiting visual response to all categories of images, but selectivity for human faces. C) Rasters and spike density functions of example neurons exhibiting pre-saccadic and post-saccadic activity. D) Top panel, task schematic. Middle panel depicts a single neuron’s response in different contexts. Bottom panel shows a single neuron’s response to different call types in different contexts.

### Neurons in Most lPFC Subregions Respond During A WM Task

The relationship between the observed responses in all epochs of the DML task and the lPFC subregions in which they were recorded is shown in Figure 2A, which presents the proportion of all units recorded at a given array location that were significantly modulated within each task epoch, pooled across all sessions in which the DML task was run. Overall, we found that single neurons exhibited significant modulations in all task epochs across all lPFC subregions, but noted that the proportion of units with delay activity was relatively lower in areas 9 and 10. Detailed analyses of these data have been reported previously (Wong et al. 2023).

**Figure 2.**
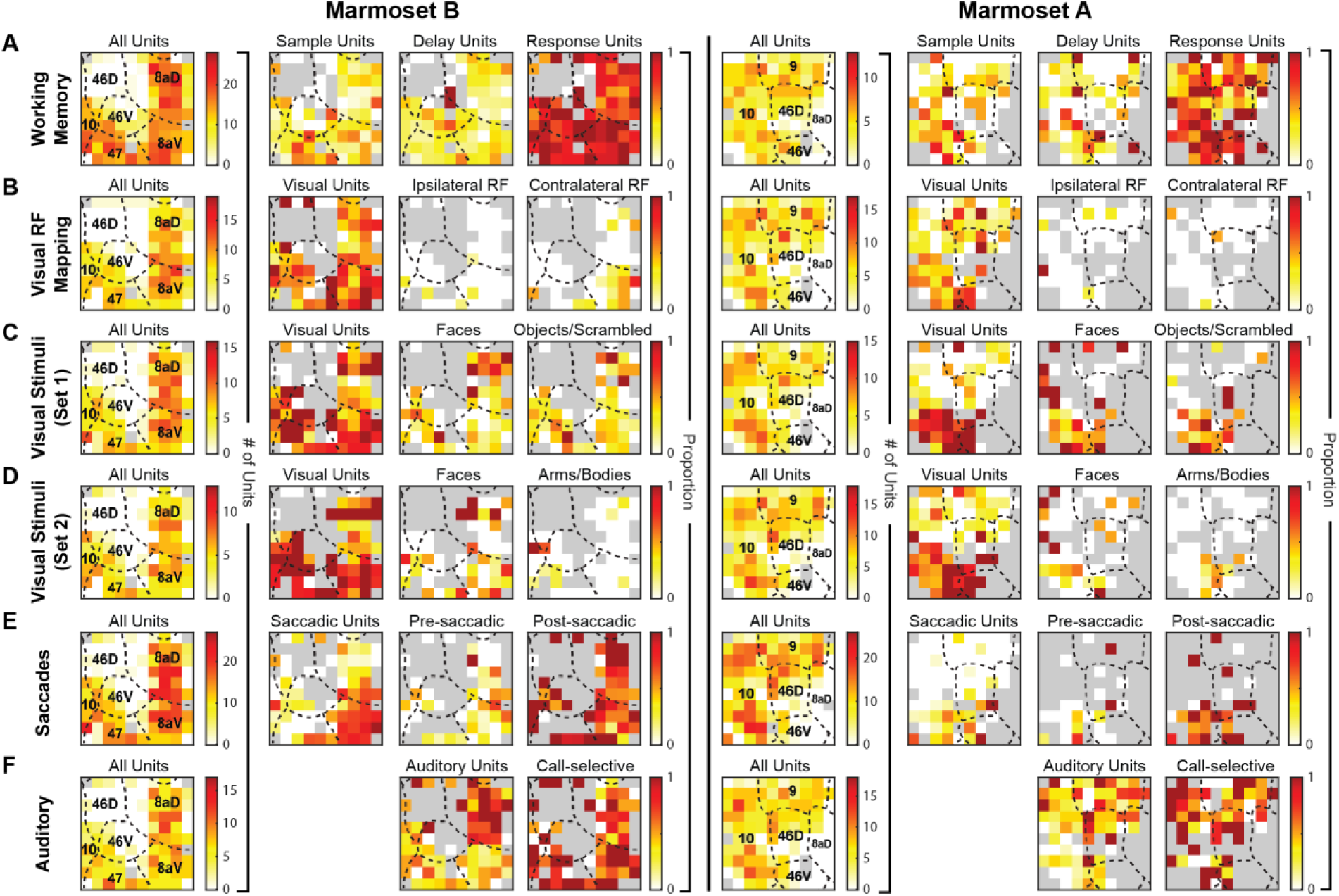
Distribution of task-modulated units and units responsive to different stimulus modalities. Array locations were reconstructed using high-resolution MRIs and superimposed on a standardized marmoset brain, area boundaries from Paxinos et al. (2012). The first column (left column) represents the total number of units found across sessions and its distribution on the array. Second column represents the proportion of units compared to the first column. The third and/or fourth column represents the proportion of units compared to the second column. Grey depicts array locations at which well-isolated single units were not observed.

### PFC Visual Receptive Fields were Largely Bilateral

We carried out separate two-way ANOVAs on discharge rates of lPFC neurons with factors of epoch (baseline – 0-200ms prior to stimulus onset, visual 50-150ms following stimulus onset) and location (contralateral, ipsilateral), to determine rough estimates of lateralization of their visual receptive fields. Neurons exhibiting a significant interaction and/or a main effect of epoch were considered visual. In Marmoset B, 225/416 (54.1%) neurons were classified as visual, of which 34 and 6 neurons had contralateral and ipsilateral receptive fields, respectively. In Marmoset A, 108/365 (37.0%) neurons were classified as visual, with 3 and 5 neurons having well-defined contralateral and ipsilateral receptive fields, respectively. Overall, most neurons with visual activity exhibited these responses across a broad range of locations encompassing both ipsi- and contralateral visual fields. The proportions of all recorded units exhibiting statistically significant visual responses and location selective responses across lPFC subregions is depicted in Figure 2B. We found units with visual activity at all locations at which we found well isolated single units, with a notable cluster of neurons in 8aV that had contralateral receptive fields.

### lPFC Neurons Respond to Many Categories of Visual Stimuli

In separate recording sessions, we presented marmosets with two distinct sets of categorical images. Set one included images of human faces, marmoset faces, scrambled faces, and objects. Set two comprised images of marmoset arms, bodies, and faces. To assess whether marmoset lPFC neurons responded to these categorical images, we performed two-way ANOVAs for both the baseline and post-stimulus onset epochs. Neurons demonstrating a significant interaction and/or a main effect of epoch were identified as visually responsive.

For the sessions presenting the first set of images, the response patterns were as follows: In Marmoset B, 61.4% (234/381) of neurons showed visual activity, with 26.9% responsive to human faces, 18.8% to marmoset faces, and 31.2% to objects/scrambled images. In Marmoset A, 37.0% (118/319) exhibited visual activity, with respective responsiveness of 33.1% to human faces, 33.1% to marmoset faces, and 39.8% to objects/scrambled images. Notably, neurons responsive to faces showed higher discharge rates for human faces than for marmoset faces (one-sample t-test: p < .001).

In sessions with the second set of images (marmoset arms, bodies, and faces), Marmoset B had 68.2% (180/264) of neurons displaying visual activity, with 31.1% responding to faces and 25.6% to arms/bodies. In Marmoset A, 37.2% (148/398) showed visual activity, with 26.4% responsive to faces and 19.6% to arms/bodies.

To quantify differences in activity between categories and individual images, we calculated a selectivity index. Supplementary Figure 2 shows the averaged selectivity indices for all responsive neurons at each array location, revealing a tendency for marmoset PFC neurons to prefer individual images over specific categories. We observed visual activity across all locations with well-isolated single units (Figure 2 C-D) and noted clusters of face-selective neurons in areas 8aD, 8aV, and area 10 (Figure 2 C-D).

### Saccade-Related Responses in Marmoset lPFC Neurons

To determine whether marmoset lPFC neurons exhibited saccade-related activity, we carried out separate one-way ANOVAs at each epoch (baseline, pre-saccadic, peri-saccadic and post-saccadic). Neurons with a main effect were considered saccade-related. In Marmoset B, 314/645 (48.7%) neurons were saccade-related, with 12.4% of neurons having pre-saccadic, 30.2% having peri-saccadic and 34.9% having post-saccadic activity. In Marmoset A, 70/717 (9.8%) neurons were saccade-related, with 2.4%, 5.2% and 7.0% of neurons having pre-, peri-and post-saccadic activity, respectively. In general, many PFC neurons had activity related to spontaneous saccades with a bias toward post-saccadic responses. Saccade-related units were found to be distributed throughout the PFC, with clusters of neurons in areas 8aD, 8aV, 47, 46V and dorsal area 10 (Figure 2E).

### lPFC Neurons are Responsive to but not Selective for Conspecific Calls

To determine whether marmoset PFC neurons exhibited responses to auditory stimuli consisting of conspecific calls, we carried out three-way mixed ANOVAs for the analysis of call-type (phee, twitter, trill, trillphee, chirp, ek, chatter, tsik, scrambled) at each epoch (before and after auditory stimuli) for each experimental context (head-restrained in chair, head-unrestrained in chair and head-unrestrained in home cage). Neurons with a significant interaction and/or a main effect of epoch were considered as having auditory responses.

In Marmoset B, 191/373 (51.2%) neurons had auditory activity, 84 (22.5%) neurons had a three-way interaction, 46 (12.3%) neurons had an interaction between call type and epoch, 44 (11.7%) neurons had an interaction between context and epoch, and 41 (11.0%) neurons had a main effect of epoch. Of the 67 neurons that had an interaction of call type and epoch (including 21 which had a higher order interaction), 14 were responsive to one call type, 9 were responsive to two call types, 5 were responsive to three call types and 39 were responsive to more than three call types (excluding scrambled). In Marmoset A, 157/418 (37.6%) neurons had auditory activity: 65 (15.5%) neurons had a three-way interaction, 36 (8.6%) neurons had an interaction between call type and epoch, 17 (4.1%) neurons had an interaction between context and epoch, and 49 (11.7%) neurons had a main effect of epoch. Of the 51 neurons that had an interaction of call type and epoch (including 15 which had a higher order interaction), 13 were responsive to one call type, 16 were responsive to two call types, 7 were responsive to three call types and 15 were responsive to more than three call types (excluding scrambled). In neurons with a 3-way interaction, we examined conducted Bonferroni corrected pairwise comparisons to examine differences in discharge activity from baseline for different call types in different contexts. We observed no systematic effect of experimental context on responses to different call types; a neuron’s response profile for calls in one context did not predict responses in another context. Altogether, we found that most call-responsive neurons exhibited responses to multiple call types. Auditory and call-selective neurons were observed throughout all PFC subregions, with a higher proportion of auditory neurons in areas 8aD, 46D, 9 and the dorsal portion of area 10 (Figure 2F).

### Natural Grouping of Neural Responses Reveals Spatially Segregated Clusters in Marmoset lPFC

We used unsupervised algorithms (Kiani et al. 2015) to identify natural groupings of neurons based on their response covariation within recording sessions. With these objectively identified groupings of neurons, we projected back onto the arrays to determine whether neurons were spatially segregated in a topographic manner. Though lPFC regions may have similar response properties, this method can reveal a topography that is defined at the population level. Figure 3 shows the MDS-filtered dissimilarity matrix colour map in which similar colours depict neurons with similar responses. (see “Natural Grouping of Neural Responses” in Methods).

**Figure 3.**
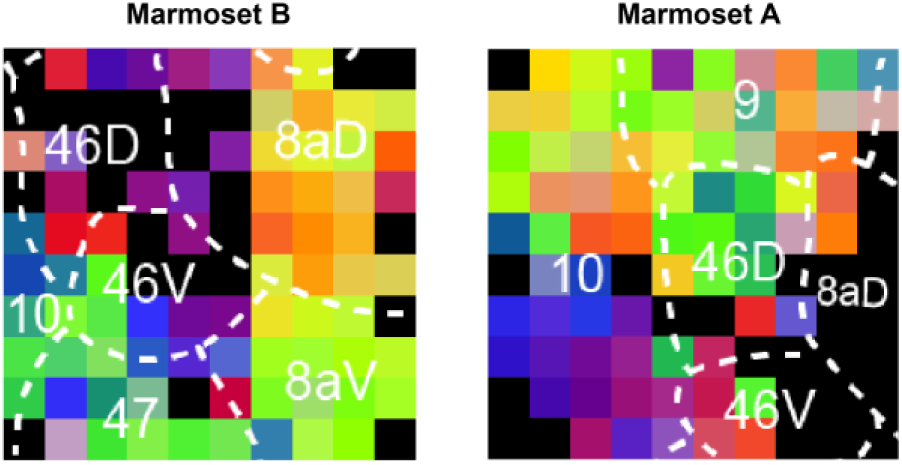
Natural grouping of neural responses reveals spatially segregated clusters. For each session, we identified natural physiological groupings of recorded units based on the dissimilarity of their responses (see methods). To create a low dimensional representation and to explore the spatial relationship between recording channels, we chose a unique colour for each recording channel based on its location in the 2-dimensional MDS map and a 2D colour map. Locations on the array with similar colours represent natural physiological groupings of recorded units.

Based on this analysis, we found that that the PFC was divided into functional subregions, similar to the cytoarchitectural boundaries we outlined, and potentially into further smaller sub-regions in area 10, dorsal and ventral. Overall, this indicates that although single PFC neurons across PFC subregions are responsive to many stimuli and modulated during a WM task, responses within a given subregion are more similar to each other than those across subregions. This is suggestive of a degree of functional localization in marmoset PFC.

## Discussion

The common marmoset is a model of growing popularity for studies of cognitive processes and has great potential as a model animal for investigations of the neural basis of social cognition (Samandra et al. 2022). The role of the PFC in cognitive processes and the specific roles of lPFC subregions have been investigated in myriad neurophysiological studies in rhesus macaques (Hoshi, 2006; Wise, 2008), however far fewer studies have investigated the responses of marmoset lPFC neurons to social and non-social visual and auditory stimuli, or during cognitive tasks. Here, we sought to characterize the responses of marmoset lPFC neurons across a suite of subregions by employing a broad test battery including a spatial working memory task, visual receptive field mapping task, the presentation of differing categories of social and non-social images, social and non-social auditory stimuli. Task-modulated neurons and neurons responsive to different stimulus modalities were found throughout the lPFC of marmosets, with relatively subtle variations across recording locations and hence cytoarchitectonic boundaries (see summaries in Table 1 and Figure 4). Interestingly, in spite of the broad distribution of responses to a widely varied suite of visual and auditory stimuli, we found that the recorded population of neurons in the PFC was not homogeneous but could be divided into smaller subregions with a clustering, similar to cytoarchitectural boundaries based on task-independent covariation of neural responses. This suggests the responsiveness of lPFC neurons across areal boundaries during many task epochs and to many stimuli is superimposed onto a basic pattern of greater discharge covariation within rather than between lPFC subregions.

**Table 1.** Summary of lPFC subregions within which task-modulated units and units responsive to varying stimulus modalities were found.

|  | Cortical Area |  |  |  |  |  |  |
| --- | --- | --- | --- | --- | --- | --- | --- |
| Modality | 8aV | 8aD | 9 | 10 | 46D | 46V | 47 |
| WM: Sample Units | ✓ | ✓ | ✓ | ✓ | ✓ | ✓ | ✓ |
| WM: Delay Units | ✓ | ✓ | ✓ | ✓ | ✓ | ✓ | ✓ |
| WM: Response Units | ✓ | ✓ | ✓ | ✓ | ✓ | ✓ | ✓ |
| Visual Units | ✓ | ✓ | ✓ | ✓ | ✓ | ✓ | ✓ |
| Contralateral RF | ✓ | ✓ |  |  |  |  |  |
| Faces | ✓ | ✓ |  | ✓ |  | ✓ | ✓ |
| Auditory Units | ✓ | ✓ | ✓ | ✓ | ✓ | ✓ | ✓ |
| Call-selective Units | ✓ | ✓ | ✓ | ✓ | ✓ | ✓ | ✓ |
| Saccadic Units | ✓ | ✓ |  | ✓ |  | ✓ | ✓ |

**Figure 4.**
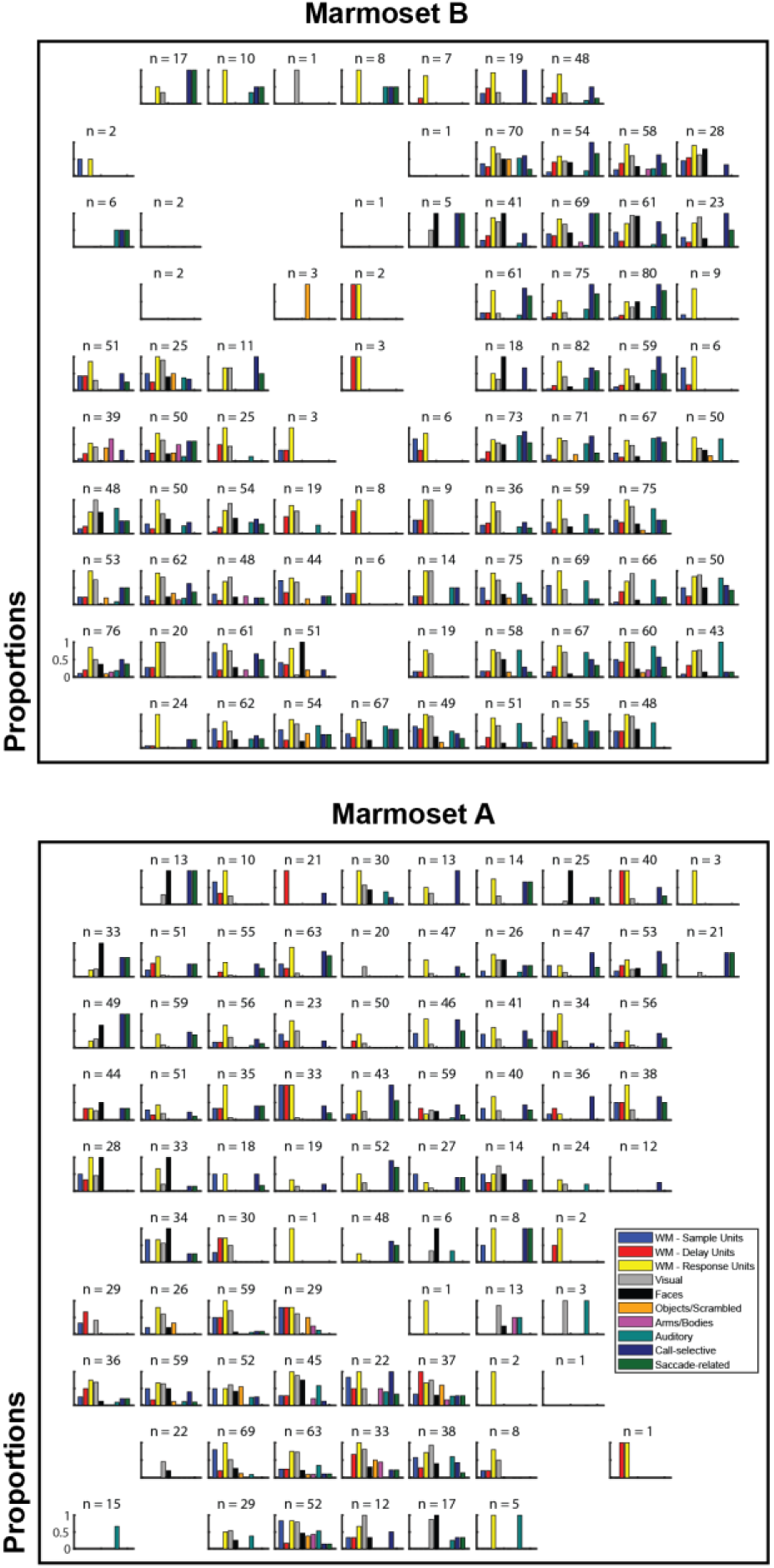
Proportions of task-modulated units and units responsive to different stimulus modalities at each electrode contact in Marmoset B (A) and Marmoset A (B). n indicates the overall total number of units at each electrode contact. Proportions were not calculated with respect to overall total number of units, rather in respect to the total number of units found in task-specific sessions.

As we have reported previously (Wong et al. 2023), we observed activity in a spatial WM task in marmoset lPFC neurons which was remarkably similar to initial reports by Fuster and Kubota in the macaque (Fuster and Alexander 1971; Kubota and Niki 1971). Like macaque lPFC neurons, in the DML task, marmoset lPFC neurons exhibited sample-, delay-, and response-related activity. We observed task-related activity in all subregions, but noted that the proportion of units with delay-related activity was relatively lower in areas 9 and 10. Given the anatomical connections of these areas to temporal cortical areas associated with higher order auditory and visual stimuli (see Burman et al. 2006), and our observations here of predominantly auditory responses in these areas, this finding suggests that these areas may not have a significant role in the short-term retention of visual information. Excitotoxic lesions of area 9 in marmosets have been shown to impair inhibitory control in a complex set-shifting task (Dias et al. 1996), suggesting that this area may be involved in more attentional shifting rather than retention, though further investigations comparing these types of tasks will be required to establish definitively the roles of areas 9 and 10 in cognitive processes.

In light of previous work establishing visual responses and visual receptive fields in lPFC neurons (Boch and Goldberg 1989) and the well-established role of lPFC in visual attention (Rossi et al. 2009), we investigated visual receptive fields of lPFC neurons. Consistent with a previous report which found robust visual responses in a broad region of PFC including areas 6DR, 8aD, 8aV, 8C and 46V (Feizpour et al. 2021), we observed units with visual responses in virtually all lPFC areas which we recorded, including areas 9, 10, and 46D, which were not sampled in that study due the locations of their array implantations which also included areas 8C, 6Va, and 6DC and importantly established visual responses in these areas as well. As in that study, we noted that the greatest proportion of visual units was found in area 8aV, and that neurons in this area exhibited contralateral response fields. This, together with our finding of a large proportion of neurons with saccade-related activity in this area, is consistent with previous evidence suggesting that these areas contain the marmoset FEF (Selvanayagam et al. 2019; Feizpour et al. 2021).

During sessions in which marmosets were presented with visual stimuli belonging to different categories, neurons with visual responses were found in all lPFC subregions. We found clusters of face-selective neurons in areas 8aV, 8aD, 10, 46V and 47, but no specific topography for arms, bodies, objects or scrambled images. A previous fMRI study in awake marmosets observed activation in frontal areas, predominately 8aV and 45, for images of marmoset faces, bodies, objects and scrambled versions of these images (Hung et al. 2015). In a subsequent fMRI study, videos containing conspecific faces evoked activity across a broad network that was similar, but stronger than the topology that was elicited using scrambled versions of the videos (Schaeffer et al. 2020). They proposed that the lateral prefrontal patches they observed, with activation peaks in areas 45 and 47, were likely face selective, while the activation extending into 8aV may have been more related to saccades made by the animals while viewing the videos. In our study, we found that areas in which neurons with face-selective responses were found overlapped with areas that are saccade-related, with neurons being both visual and saccade-related. Altogether, at the single-neuron level, we observed more wide-spread responses in the lPFC of marmosets than previously observed in fMRI studies.

We also observed robust auditory responses in lPFC neurons when marmosets were presented with sets of acoustic stimuli. Neurons with auditory responses were found throughout all lPFC subregions sampled, with the highest proportions present in areas 8aD, 9, 46D and the dorsomedial portion of area 10. In a prior study exploring auditory responses in the marmoset lPFC, neurons that were responsive to long-distance “phee” calls were identified in areas 8aV, 8aD, and 45 (Jovanovic et al. 2022). Our findings suggest a broader distribution of auditory responses across lPFC subregions, potentially due to the presentation of multiple types of calls. However, it’s noteworthy that even the solitary presentation of phee calls has been associated with increased cFos expression spanning a considerable expanse of the ventral and posterior PFC (Miller et al. 2010). We observed that neurons selective for at least one type of call were present throughout all lPFC subregions where auditory responses occurred. This finding is somewhat distinct from those in macaques, where auditory-responsive cells are predominantly localized in the vlPFC (areas 12 and 45; Romanski and Goldman-Rakic 2002).

We additionally investigated the effects of experimental context on responses of PFC neurons to marmoset calls. We found no evidence for cross-context consistency in the responsiveness of neurons, suggesting that PFC neurons may encode various combinations of call type and context rather than selectivity for a particular call type, which is then modulated by the context in which it is encountered. This is broadly consistent with the observed responses of lPFC neurons in macaques performing visual cognitive tasks, in which responses to cues are dependent upon both their identity and the task context in which they are encountered (White and Wise 1999). Our findings mirror those of Jovanovic et al. (2022) who found that neural responses to vocalizations were highly context-specific, suggesting that neural representations of social signals in primate PFC are not static, but highly flexible and likely reflect nuances in behavioural contexts (Nummela et al. 2017). Here we focused solely on the PFC, but the effects of social context on primate neocortical function are more widespread (Sliwa and Freiwald 2017; Ainsworth et al. 2021; Cléry et al. 2021).

In addition to investigating the functional properties of neurons across a suite of lPFC regions, we exploited the advantage of having simultaneously recorded units to investigate the functional connectivity of units within and across PFC regions. We used an approach devised by Kiani and colleagues (2015) in which dissimilarity indices are computed across array locations from the discharge rates of single units across the entire duration of a recording session. These values of theses indices reflect co-variations of responses that are independent of task or stimulus-related responses, are based primarily on fluctuations in correlated noise between groups, and have been proposed to reflect anatomical connectivity between clusters of neurons (Kiani et al. 2015). We additionally projected these values from the recording locations on our array onto the estimated cytoarchitectonic boundaries of the PFC areas as delineated in the atlas of Paxinos (2012). Similar to their findings in macaque, we observed functional clusters of units which corresponded broadly to the areal subdivisions of the marmoset lPFC, with some subclusters of units within these divisions. This finding is consistent with the known dominance of interareal anatomical connections within lPFC subregions (Reser et al. 2013), as well as the observation of “patchy” connectivity across lPFC columns (Watakabe et al. 2023). Altogether, our data suggest a spatial segregation of anatomically connected neurons within marmoset lPFC which may correspond to anatomically defined lPFC subregions. It should be noted however that cytoarchitectural boundaries are notoriously difficult to determine in the relatively undifferentiated lPFC of the marmoset (Reser et al. 2013) so a direct correspondence between dissimilarity values and cytoarchitecture is best interpreted with some caution. Nonetheless, these data do provide evidence of task-independent spatial clustering of neurons in marmoset lPFC.

The quest to understand how anatomical subdivisions of the lPFC relate to sensory, motor, and cognitive functions is a long-standing pursuit in cognitive neurosciences, as outlined by Wilson et al. (2010). Our observations, similar to numerous electrophysiological studies conducted in rhesus macaques, encompass task-based responses, reactions to a wide array of social and non-social stimuli across both auditory and visual modalities, and discharge rate variations contingent on the experimental context (refer to Wise, 2008 for a review). Generally, these modulations in discharge rates are distributed across multiple prefrontal areas. However, exceptions include more localized face-selective responses in areas 47 and 8aV, and contralateral visual receptive fields and saccade-related responses in area 8aV. This latter region is likely the marmoset homologue of the marmoset FEF (Selvanayagam et al. 2019; Feizpour et al. 2021).

This too is consistent with observations in macaque (Kiani et al. 2015). Overall, these similarities in organization to macaque PFC and the distributed nature of many social and non-social representations in marmoset PFC provide supporting evidence for the validity of the marmoset model in investigations of cognitive control and social cognition. An intriguing possibility supported by our data here is that, as in humans and macaques, the marmoset PFC as a whole is a crucial part of a multi-demand system engaged to deal with diverse cognitive demands (Duncan 2010), of which social interactions and vocal communication would be a major part. As has been noted previously these situations require the integration of diverse sources of visual, auditory, context-based information, and the evaluation of outcomes in the service of behavioural goals (Jovanovic et al. 2022; Samandra et al. 2022; Grijseels et al. 2023). More naturalistic studies combining social interactions and vocal communication with neural recordings, so uniquely suited to the marmoset model, have the potential to advance our understanding of both PFC function in general and its specific role in social cognition.

## Acknowledgements

The authors wish to thank Cheryl Vander Tuin, Whitney Froese, and Hannah Pettypiece for animal preparation and care and Peter Zeman for technical assistance. We would also like to thank David Everling for assistance with touchscreen testing. This research was supported by the Canadian Institutes of Health Research (CIHR) grant FRN148365 to SE and the Canada First Research Excellence Fund to BrainsCAN. RW was also supported by the Canada First Research Excellence Fund to BrainsCAN and the Next Generation Networks for Neuroscience (NeuroNex). JS was supported by a Natural Sciences and Engineering Research Council (NSERC) Canadian Graduate Scholarship (Doctoral).

## Author Contributions

RW performed experiments, analysed data, prepared figures and wrote the manuscript. JS performed experiments and assisted in data analysis. KJ assisted in surgeries and data analysis. SE designed experiments and performed surgeries. All authors edited the manuscript and SE approved the final version.

---

**Supplementary Figure 1.**
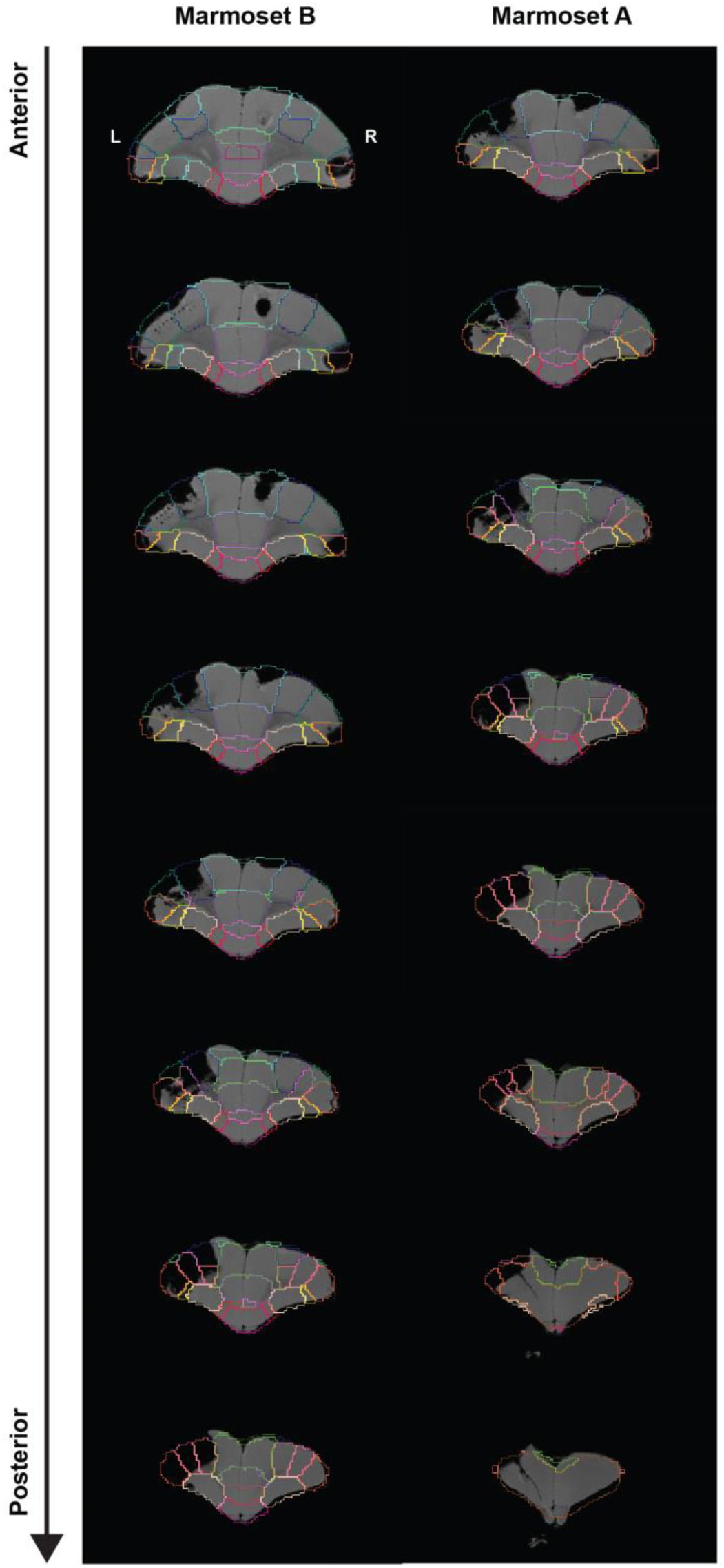
Reconstruction of array implantation sites. T2-weighted MRIs were acquired and nonlinearly registered to the NIH template brain that contains the location of cytoarchitectonic boundaries of the Paxinos atlas (see methods). Each consecutive coronal slice (anterior to posterior) is spaced by 400 µm.

**Supplementary Figure 2.**
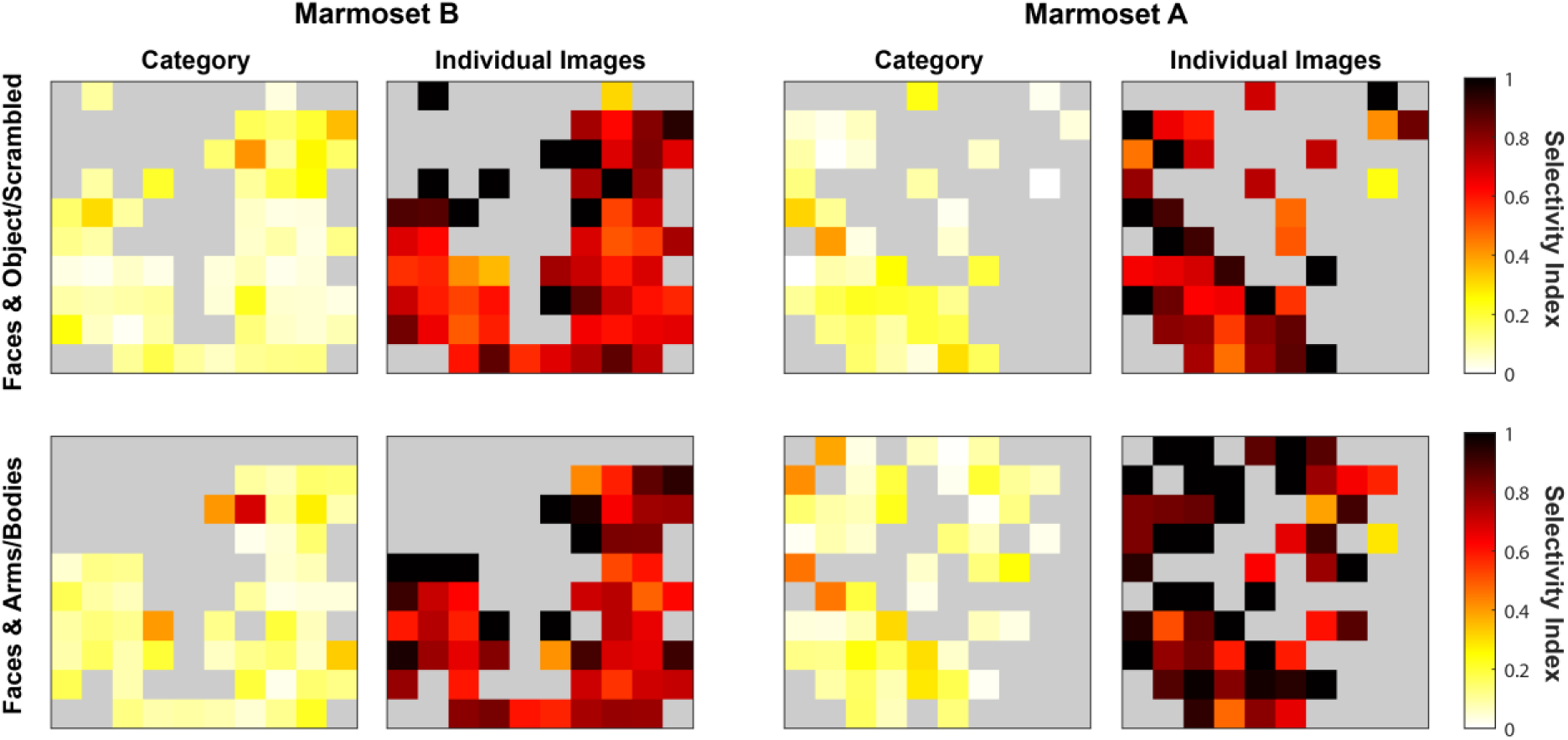
Averaged selectivity indices computed for all responsive neurons at each array location for between categories of images, as well as between individual images.

## Notes

### Competing Interest Statement

The authors have declared no competing interest.

